# Has AlphaFold 3 reached its success for RNAs?

**DOI:** 10.1101/2024.06.13.598780

**Authors:** Clément Bernard, Guillaume Postic, Sahar Ghannay, Fariza Tahi

## Abstract

Predicting the 3D structure of RNA is a significant challenge despite ongoing advancements in the field. Although AlphaFold has successfully addressed this problem for proteins, RNA structure prediction raises difficulties due to fundamental differences between proteins and RNAs, which hinder direct adaptation. The latest release of AlphaFold, AlphaFold 3, has broadened its scope to include multiple different molecules like DNA, ligands and RNA. While the article discusses the results of the last CASP-RNA dataset, the scope of performances and the limitations for RNAs are unclear. In this article, we provide a comprehensive analysis of the performances of AlphaFold 3 in the prediction of RNA 3D structures. Through an extensive benchmark over five different test sets, we discuss the performances and limitations of AlphaFold 3. We also compare its performances with ten existing state-of-the-art *ab initio*, template-based and deep-learning approaches. Our results are freely available on the EvryRNA platform: https://evryrna.ibisc.univ-evry.fr/evryrna/alphafold3/.

## Introduction

Ribonucleic acids (RNAs) are fundamental molecules crucial to cellular activities. While their functions are directly linked to their structures, the prediction of the latter remains an open challenge to be addressed. Knowing the structure of RNA could be of great interest for drug design or the comprehension of biological processes like cancer (1). While experimental methods like X-ray crystallography, NMR, or cryo-EM can determine the RNA 3D structures, their use is costly (in terms of time and resources) and hardly scalable for the number of RNAs in the living. Computational approaches have emerged with *ab initio*, template-based and, more recently, deep learning methods. *Ab initio* methods (2–11) tend to reproduce the physics of the system, with force field applied to a coarse-grained representation (low-resolution where a nucleotide is replaced by some of its atoms). The template-based approaches (12–21) create a mapping between sequences and fragments of structure before refining the assembled structures.

With the recent success of AlphaFold for proteins (22, 23), approaches have been made to replicate its success to RNAs. Directly using protein methods to infer RNA 3D structures is impossible, as RNAs and proteins are biologically different molecules. Current methods try to adapt what exists for proteins to RNAs like DeepFoldRNA (24), RhoFold (25), DrFold (26), NuFold (27), and trRosettaRNA (28). They consider coarse-grained representation and predict Euclidean transformation before reconstructing the fullatom structure. The use of torsional angles is also adapted to RNAs, with either the standard torsional angles (25, 27) or angles from their coarse-grained representations (24, 26).

While being better than existing template-based or *ab initio* methods, deep learning approaches do not solve the prediction of RNA structures yet, as shown in CASP-RNA (29) and in our recent benchmark State-of-the-RNArt (30). Recently, a critical review (31) explains the reasons why the AlphaFold for RNA has not yet happened, and might not arrive for the next decades. However, AlphaFold has released its latest version, named AlphaFold 3 (32), that extends its predictions to different molecules, including RNAs. In this work, we aim to provide a response to (31) in order to know if AlphaFold 3 meets its success for RNAs.

To extend its range of molecules, AlphaFold 3 has made changes in its architecture to better adapt to the variety of available inputs. It no longer relies on torsional angles to prevent the restriction to specific molecules, as was the case in AlphaFold 2 (23). It directly predicts atom coordinates with the use of a multi-cross diffusion model. Through a benchmark on CASP-RNA (29), the authors mentioned good results, but AlphaFold 3 did not outperform human-helped methods. Furthermore, it is not clear what the current limitations are and how well it performs compared to state-of-the-art solutions.

This article aims to provide a comprehensive extension on the evaluation and benchmark of AlphaFold 3 for RNAs. We first describe the main differences between RNAs and proteins to highlight the challenges of RNA 3D structure prediction and we describe the AlphaFold 3 solution before discussing the benchmark we did. Then, we evaluate AlphaFold 3 and comment on the results of AlphaFold 3 and the current limitations of the model. Our benchmark also compares the performances with state-of-the-art solutions to provide a complete comparison. The results and the data are freely available and usable in the EvryRNA platform: https://evryrna.ibisc.univ-evry.fr/evryrna/alphafold3/.

### RNAs vs proteins

RNAs and proteins are both molecules that play crucial roles in the living. They share the characteristic of having a 3D structure that directly defines their function. This section discusses the differences between RNAs and proteins, high-lighting the reasons why adapting existing protein models has been challenging.

RNAs comprise four nucleotides (A, C, G and U), whereas proteins comprise 20 amino acids. This difference has a high consequence on the adaptation of protein algorithms to RNA. The vocabulary available for RNA is limited to four unique elements, making the use of protein vocabulary not directly adaptable. The sequence length of RNA molecules also has a high variability (from a dozen to thousands of nucleotides) compared to proteins (around a hundred of amino acids).

A major difference between RNAs and proteins lies in the folding stabilisation. RNA structure is maintained by base pairing and base stacking, while protein structure is supported by hydrogen interactions in the skeleton. The protein backbone is also modelled by torsion angles (Φ and Ψ) for each amino acid because the peptide bond is planar. This is not the case for RNA, where each nucleotide can be described by six torsion angles (*α, β, γ, δ, ϵ, ξ*) and the sugar-pucker conformation (*χ*). An approximation usually involves pseudo-torsion *η* and *θ* (33). Protein models learn a conformational mechanism fundamentally different from the RNA folding process, where adaptations should be made for RNAs (34).

The nature of pairwise interactions of RNA 3D molecules differ from those of proteins. The RNA interactions can be made through three different edges of the RNA base: WC edge, Hoogsten edge and sugar edge shown in Figure 1. In addition, the orientation of the glycosidic bonds gives another property to an interaction: *cis* or *trans*. The combination of edge and orientation gives 12 possibilities of interaction between bases. The standard Watson-Crick (WC) base pair corresponds to the cis WC/WC pairing. Given the orientations (*cis* or *trans*), the edges and the base pairing, there are more than 200 possible base pairs. Only the standard WC pairs (cis WC/WC) of AU and CG (and also GU wobble pair) are used for the 2D structure representation. RNA bases also have common patterns of interactions, where a base stacks on another one. The base-stacking (35, 36) refers to the four base-stacking types from relative orientations (upward, downward, outward and inward) (37). The extended secondary and tertiary interactions (long-range base pairs) play a crucial role in the overall topology of the RNA folding process. They help stabilise the structure and can not be ignored when working on RNA 3D structures.

**Figure 1.**
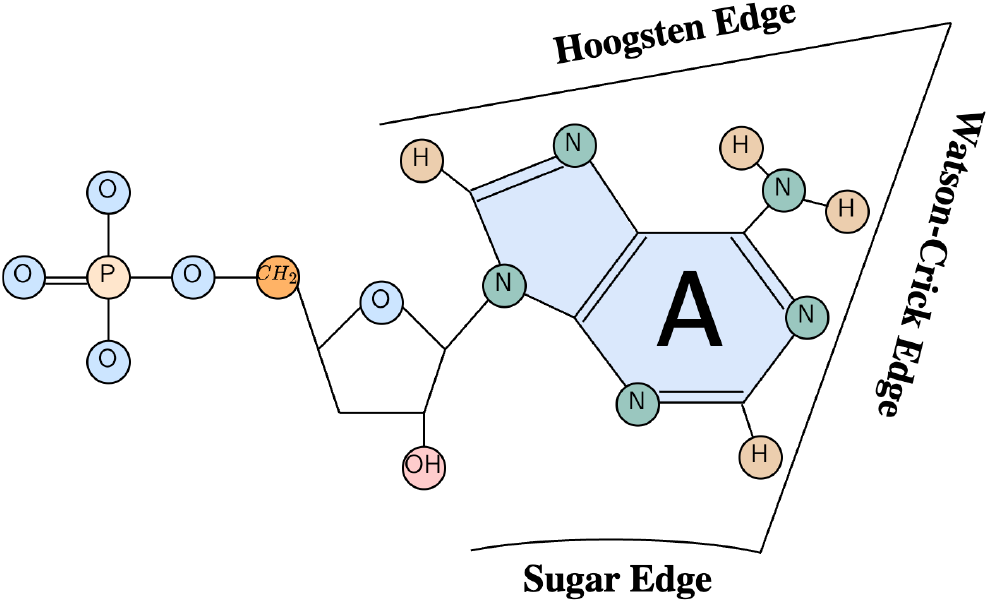
Description of the three different edges of the Adenine RNA nucleotide: Watson-Crick edge, Hoogsten edge and Sugar edge. The three other nucleotides share similar edges.

The stability of the RNA and protein structures is different. Proteins tend to have a stable structure corresponding to a minimum in the energy landscape. On the contrary, RNAs can have multiple conformations depending on the environmental condition. Those foldings are of equivalent validity as they are all local minimums in the energy landscape. Ideally, finding an RNA 3D structure means finding the native foldings of RNA, and should output multiple structures with confidence scores. However, the available structures found in the PDB do not provide access to this diversity of structures.

There is a huge disparity in the protein and RNA data. Even if there is a higher proportion of RNAs than proteins in the living, this is not reflected in the available data: only a small amount of 3D RNA structures are known. Up to June 2024, 7,759 RNA structures were deposited in the Protein Data Bank (38), compared to 216,212 protein structures. The quality and diversity of data are also different: a huge proportion of RNAs come from the same families. It implies several redundant structures that could prevent a model from being generalized to other families. In addition, a huge amount of RNA families have not yet solved structures in the PDB, usually long RNAs. This means there is no balanced and representative proportion of RNA families through the known structures.

Finally, no standard dataset has been used through the community for RNAs. Each research group uses its dataset with different preprocessing associated. It prevents using deep learning methods, as a lot of work is needed for a clean dataset. While the community agrees to use RNA-Puzzles (39–43) or the newly CASP-RNA (29) to test the generalization of proposed models, no clear training set is available. The first solution has been RNANet (44), developed in our lab to solve this issue. It is a database that uses MySQL and gathers diverse RNA information to train deep learning methods. A new approach, RNA3DB (45), creates independent datasets for deep learning approaches where clustering is done based on sequence and structure disparity.

### AlphaFold 3

Building on the recent success of AlphaFold 2 (23) in protein structure prediction, AlphaFold 3 (32) expands its predictions to structures from all molecules available in the PDB (38). The new release directly predicts the raw coordinates of atoms, eliminating the specificities associated with the AlphaFold 2 architecture. The authors highlight several differences from the previous architecture, contributing to successful predictions of a wide range of molecules. One key difference is the introduction of a diffusion model that reconstructs coordinates from the residue level to the atom level. In the case of RNA, AlphaFold 3 has been evaluated on the CASP-RNA dataset (29), demonstrating improved predictions compared to RosettaFold2NA (46) and AIchemy_RNA (25) (the best AI-based submission in the competition). Despite these advancements, AlphaFold 3’s performance lags behind AIchemy_RNA2 (47) (the top human-expert-aided submission).

In this section we present briefly the AlphaFold 3 architecture, the training procedure, the differences from previous approaches and the limitations mentioned in the article.

### AlphaFold 3 architecture

AlphaFold 3 takes as inputs a sequence of a given molecule (amino acids for proteins, nucleotides for RNAs, etc) and embeds it to different main blocks: the input embedder, the pairformer and the diffusion module. The sequence input is represented as tokens, where a token is considered a nucleotide for RNAs. An MSA module is added before the Pairformer, which reduces its overall importance in the network. A different input now embeds the information, the pair representation. The architecture also adds confidence measures like pLDDT (modified local distance difference test), PAE (predicted aligned error) and PDE (distance error matrix).

A novelty brought by the third version of AlphaFold 3 is the replacement of the structural module with a diffusion module. This module trains a denoiser to remove Gaussian noise from coarse-grained representation (one atom per nucleotide for RNA/DNA or per amino acid for protein) to full atom representation. No explicit geometry is involved, which differs from existing recent solutions (25–28, 48). Instead, the diffusion module is trained to reconstruct 48 different versions of the reference structure (randomly rotating, translating, and noised). The training loss for the diffusion module is a weighted aligned MSE loss, which is then incremented by a structure-based loss based on smooth LDDT for the finetuning part.

The final architecture of AlphaFold 3 can predict up to 5120 tokens, for an inference time of 347 seconds on 16 NVIDIA A100 GPUs (32).

### Training procedure

AlphaFold 3 was trained using a mixture of five datasets (PDB structures, transcription factors and distilled datasets). The RNA data came from the Rfam (49), RNACentral (50), and Nucleotide collection (51). Only structures with a release date below September 2021 were used for the training, and structures with a release time between September 2021 and January 2023 were for the validation set. The RNA distilled dataset includes predictions from AlphaFold 3 on representative clusters from Rfam (49) (using a cutoff of 90% of sequence identity and 80% coverage to cluster the sequences into clusters). It leads to an overall of around 65,000 structures for RNAs.

During the training loop, one dataset is sampled, and an example is drawn. Then, a structural crop is made using different strategies: contiguous, spatial, or spatial interface cropping. This strategy was presented in AlphaFold-Multimer (52), where the idea is to tackle the high memory and computing required when dealing with long structures. Therefore, the model is trained on cropped segments of full-length molecules, where each subregion is a contiguous block of tokens extracted depending on previously mentioned strategies.

There are four main stages of training for AlphaFold 3. The first uses sequences cropped to 384 tokens, while the rest are considered as fine-tuning. The second stage increases the crop size to 640. The third stage has a crop size of 768 tokens, and the transcription factor distillation sets were made accessible. Finally, the last fine-tuning stage enabled the training of the PAE head and removed the structure-based losses. The initial training took ten days, the second stage 3 days, the third stage took five days, and the last fine-tuning took two days on 256 NVIDIA A100 GPUs (32).

### Differences with AlphaFold 2

AlphaFold last version had to make some changes to its previous version to be adaptable to a large variety of molecules. The different architectures are presented in Figure 2. The first difference is the input, which is not restricted to amino acids. Each residue can be one value among 32: 20 amino acids, 4 RNA nucleotides, 4 DNA nucleotides, a gap (for MSA representation) and three unknown values (one for amino acid, one for RNA nucleotide and one for DNA nucleotide). A second difference is the smaller consideration of the MSA embedding. Instead, this pair representation mainly transmits the information to the diffusion module. The MSA module is only used by the Pairformer module. The third main difference is the removal of the structure module to a diffusion module. This diffusion module does not explicitly consider equivariant processing. Finally, AlphaFold 3 directly outputs the full atom positions, contrary to backbone frames and torsional angles for AlphaFold 2. There are also small differences in the architectures, like the addition of relative token encoding or sequence-local atom attention (where the attention is restricted to a subset of 32 atoms with 128 atoms nearby in sequence space) to prevent huge computation. They also replaced the ReLU activation with SwiGLU, which has proven to be more efficient in their experiments. Each model has a different confidence head: AlphaFold 2 uses experimentally resolved score, pLDDT and pTM for confidence, masked MSA loss, FAPE, structure violation, histogram loss and side chain loss for the final global loss. AlphaFold 3 also uses experimentally resolved score, pLDDT (and pTM) and histogram losses but integrates also the PDE, the diffusion loss and the PAE.

**Figure 2.**
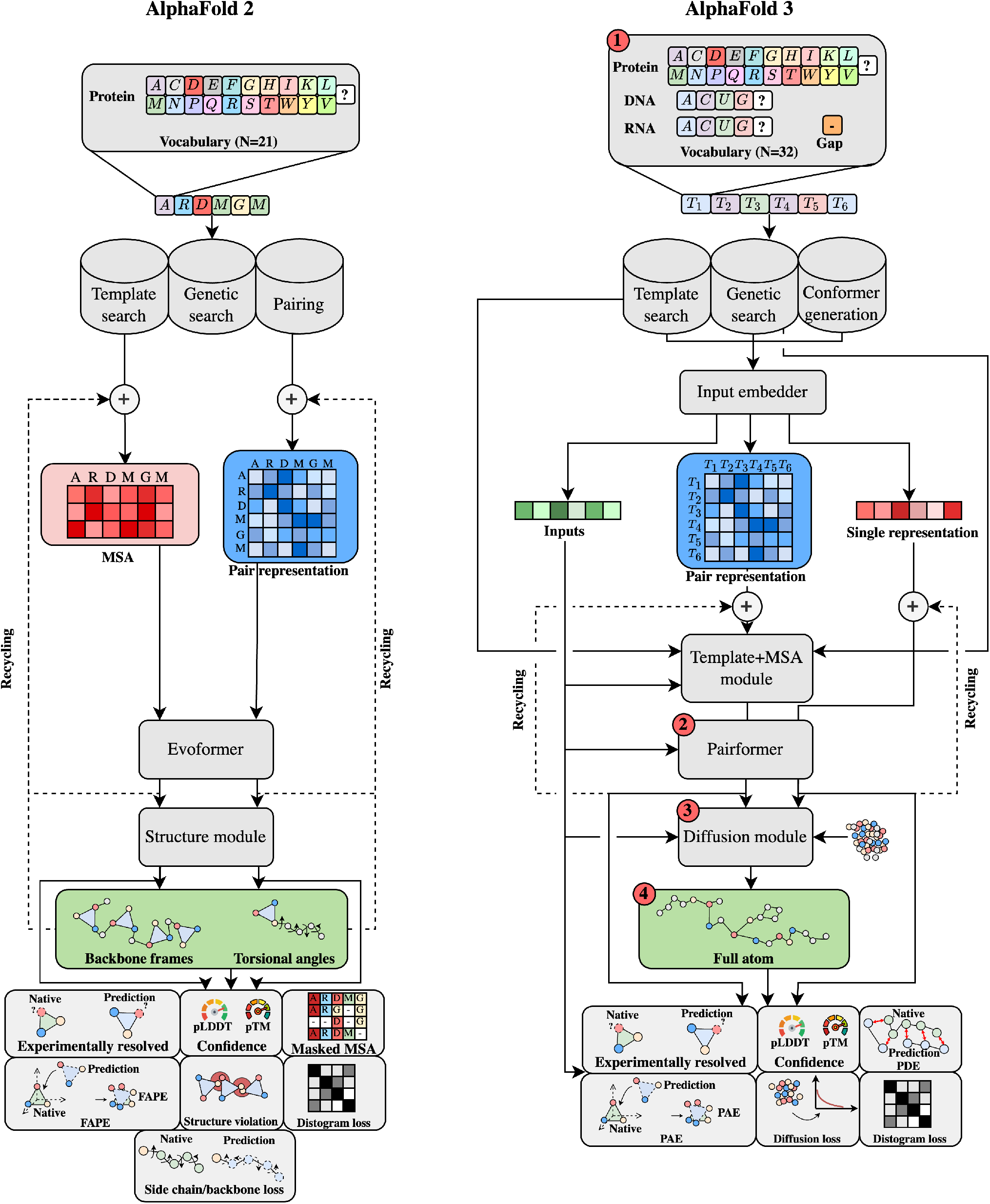
Architectures of AlphaFold 2 (23) (left) and AlphaFold 3 (32) (right). The key differences are pointed out with red numbers. 1) Difference in terms of inputs: AlphaFold 3 can predict structures of different molecules (proteins, DNAs, RNAs, ligands and ions). 2) The Evoformer is replaced by the Pairformer module, where the MSA is less important, and the pair represents more weights in the network. 3) A diffusion module replaces the structure module to output the final structure. 4) The outputs of the models are different: AlphaFold 3 now outputs directly the positions of atoms, compared to backbone frames and torsional angles for AlphaFold 2.

## Limitations

AlphaFold 3 has brought novel architecture to enable different molecular structure predictions but remains limited by the diffusion module. Indeed, diffusion-based models tend to have hallucination (53). This is reflected by plausible structures in unstructured regions, which did not happen in the previous version of AlphaFold (observed by the authors). Another limitation is the stereochemistry violations: chirality violation and the production of clashing atoms in the predictions. The chirality violation refers to the orientation and symmetry of molecules, where violation could disrupt molecular recognition, binding, and interactions with other molecules. The clashing atoms refer to atoms that are positioned too close to one another, resulting in steric clashes or overlaps. These clashes indicate that the atoms are occupying the same or very close spatial regions, which is energetically unfavourable and physically unrealistic (due to the repulsive forces between the electron clouds of the atoms). The final explicit limitation mentioned by the authors is the non-dynamical predictions, where only static structures are output. We could also mention the non-availability of the code, which prevents its large use for the community. A web server is available at https://alphafoldserver.com/ to make online predictions, but it is limited to twenty predictions per day (when writing this article).

## Benchmark

To assess the extent of AlphaFold 3 performances, we have evaluated and compared it with other state-of-the-art methods on five datasets. This section describes the datasets, the methods as well as the metrics used to evaluate AlphaFold 3.

## Datasets

To evaluate the prediction of RNA structures, we considered the following five test sets, with the first three from our previous work (30).

- **RNA-Puzzles**: the first dataset is composed of the single-stranded structures from RNA-Puzzles (39–43), a community initiative to benchmark RNA structures. We considered only single-stranded solutions to have a fair comparison between the benchmarked models. It is composed of 22 RNAs of length between 27 and 188 nt (with a mean of 83 nt).
- **CASP-RNA**: the second test set is CASP-RNA (29) structures, which is a collaboration between the CASP team and RNA-Puzzles. It is composed of 12 RNAs with wide-range sequences, from 30 to 720 nt (with a mean of 209 nt).
- **RNASolo**: the third test set is a custom test set composed of independent structures from RNAsolo (54). We downloaded the representative RNA molecules from RNAsolo (54) with a resolution below 4 Å and removed the structures with sequence identity higher than 80%. Then, we considered only the structures with a unique Rfam family ID (49), leading to 25 non-redundant RNA molecules, with a sequence between 45 and 298 nt (and a mean of 100 nt). It can not be ensured that the structures from this dataset were not used in the training set of the different models. We keep this dataset for comparison, as we already have the results for the benchmarked methods.
- **RNA3DB_0**: this dataset is composed of a non-redundant set of structurally and sequentially independent structures from RNA3DB (45). It comprises the component #0, which is composed of orphan structures that are advised to be used as a test set. These structures do not have Rfam families (49) and include synthetic RNAs, small messenger RNAs crystallized as part of larger complexes. After removing structures with sequences below ten nucleotides and sequence identity below 80% (using CD-HIT (55)), we ended up with a dataset of 224 structures from 10 to 339 nt (with a mean of 55 nt).
- **RNA3DB_Long**: the last dataset comprises long RNA structures from RNA3DB (45). We considered structures with a release date after January 2023 to avoid any structure leakage for fair comparison. We considered structures with sequences between 800 nt (800 nt being the limit from previous test sets) and 5,000 nt, as we wanted to study the performances of long RNAs. It leads to 58 structures with a sequence between 828 and 3,619 nt (with a mean of 2005 nt). It comprises 57 ribosomal RNAs and one structure of a Group II intron.

We have also ensured that all the datasets (except RNA-Puzzles and CASP-RNA) have a sequence identity below 80% to have non-redundant structures for robust evaluation.

To comprehend and detail the predictions of AlphaFold 3, we studied in detail three main interactions in the folding of RNA 3D structures: Watson-Crick (WC), non-Watson-Crick (nWC) and stacking (STACK). The distribution of these interactions is presented in Figure 3. RNA_Puzzles, CASP-RNA and RNASolo have a similar mean count of nWC interactions, while CASP-RNA has more stacking and WC interactions. RNA3DB_0 has fewer nWC interactions, and RNA3DB_Long has a significantly higher number of all types of interactions. The count of different interactions seems related to the number of nucleotides: RNA3DB_Long has structures with long sequences compared to the other datasets. RNA3DB_0, based on orphan structures, displays fewer classic RNA interactions than the other datasets. RNA3DB_0 and RNA3DB_Long have different challenges: RNA3DB_0 has complex structures with less common interactions, while RNA3DB_Long has structures with a complex number of interactions to reproduce.

**Figure 3.**
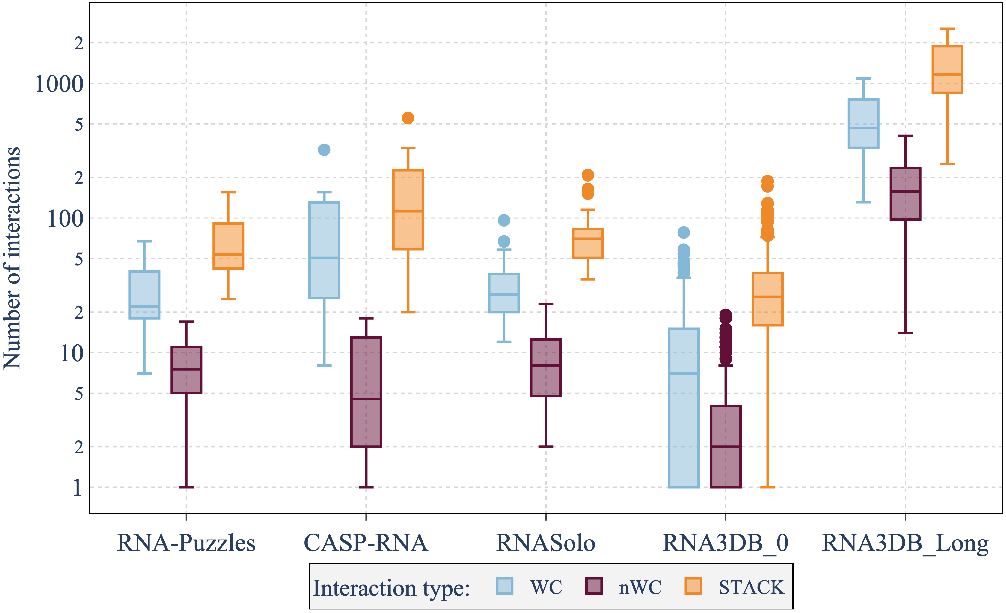
Distribution of the number of key RNA interactions for the five test sets. Interactions are either Watson-Crick (WC), non-Watson-Crick (nWC) or stacking (STACK). The logarithm scale is used as the ranges for RNA3DB_Long dataset are higher than the other test sets. Minimum values are set to 1 for the logarithm scale.

### State-of-the-art methods

Existing solutions for the prediction of RNA 3D structures are based on three main types of methods: *ab initio*, template-based and deep-learning ones. As discussed previously in our work (30), *ab initio* methods (2, 4, 6) integrate the physics of the system by usually simplifying the representation of nucleotide (coarse-grained). Instead of using all the atoms for one nucleotide, they create a low-resolution representation that simplifies the computation time while losing information. They use approaches like molecular dynamics (56) or Monte Carlo (57) to perform sampling in the conformational space and use a force field to simulate real environment conditions. On the other hand, template-based methods (12, 16– 18, 58) create a mapping between sequences and known motifs with, for instance, secondary structure trees (SSEs) before reconstructing the full structure from its subfragments. Finally, the recent methods tend to incorporate deep learning methods (24–28) by using attention-based architectures with self-distillation and recycling as done in AlphaFold 2 (23).

To compare the performances of AlphaFold 3 (32), we benchmarked ten approaches, the ones used in our previous work (30). For the *ab initio* methods, we benchmarked SimRNA (6), IsRNA1 (4) and RNAJP (2). Only RNAJP was used locally. For the template-based approaches, we benchmarked MC-Sym (58), Vfold3D (18), RNAComposer (17), 3dRNA (16) and Vfold-Pipeline (12). For the deep learning methods, we benchmark trRosettaRNA (28) and RhoFold (25). More details on each method are provided in our previous article (30). For RNA-Puzzles and CASP-RNA, we included in the benchmark the predictions from the official results of the competitions. We refer to them as “Challenge best” and correspond to different methods for each RNA. We normalised each prediction using RNA-tools (59) to have a standard format for all structures. It gives standardised names for chains, residues and atoms and removes ions and water.

We used the web servers with default parameters to compare available models fairly, where each user could reproduce our experiments. As we made most of the predictions using web servers, the predictions on RNA3DB_0 were hardly applicable to all the methods. Therefore, we benchmarked the RNA3DB_0 dataset with one method per approach (the quickest method per approach): RhoFold (25) for deep learning, RNAComposer (17) for template-based and RNAJP (2) for *ab initio*. For the RNA3DB_Long dataset, only AlphaFold 3 could predict structures with sequences up to 3000 nt. For this dataset, we only considered the predictions from AlphaFold 3.

### Evaluation metrics

To compare the predictions, we used the RNAdvisor tool (60) developed by our team, which enables the computation of a wide range of existing metrics in one command line. For the evaluation of RNA 3D structures, a general assessment of the folding of the structure can be done with either the root-mean-square deviation (RMSD) or its extension adding RNA features *ϵ*RMSD (61). Protein-inspired metrics can also be adapted to assess structure quality like the TM-score (62, 63) of the GDT-TS (64) (counts the number of superimposed atoms). There are also the CAD-score (65) (which measures the structural similarity in a contact-area difference-based function) and the lDDT (66) (which assesses the interatomic distance differences between a reference structure and a predicted one). Finally, RNA-specific metrics have been developed, like the P-VALUE (67) (which assesses the non-randomness of a given prediction). The INF-ALL (37) and DI (37) have been developed to consider RNA-specific interactions. The INF score incorporates canonical and non-canonical pairing with Watson-Crick (INF-WC), non-Watson-Crick (INF-NWC), and stacking (INF-STACK) interactions. The consideration of torsional angles has been developed with the mean of circular quantities (MCQ) (68). As discussed in (60), all these metrics are complementary and can infer different aspects of RNA 3D structure behaviour. For the rest of the article, we will discuss the RMSD, INF-ALL, lDDT, TM-score and MCQ and let the results on the other metrics in the Supplementary file. Indeed, the RMSD is the most used metric in the literature, and the INF-ALL incorporates key RNA interactions. The lDDT and TM-score allow for evaluating global conformations (widely used in AlphaFold 3), and MCQ gives the torsional deviation. We only mention all the metrics when comparing the different models to ensure a complete evaluation.

## Results

This section presents the results of AlphaFold 3 predictions on the discussed test sets. We start by comparing the results of AlphaFold 3 with existing solutions and then discuss in detail the link between the performances relative to sequence length. Next, we discuss the results of AlphaFold 3 on ribosomal structures (RNA3DB_Long dataset) and orphan structures (RNA3DB_0 dataset). Then, we discuss the results of specific RNA key interactions in detail before shedding light on the computation time.

### AlphaFold 3 compared to the state-of-the-art

We compare the predictions of ten existing methods presented above and AlphaFold 3 on our different test sets. Figure 4 presents the different normalised metrics computed for the prediction of the different models over the five test sets. We included all metrics to show the cumulative performances. The RNA3DB_Long dataset has only predictions from AlphaFold 3, which is the only method capable of processing long sequences. All the metrics are normalised by the maximum values and converted to be better where near to 1 and worst when near to 0.

**Figure 4.**
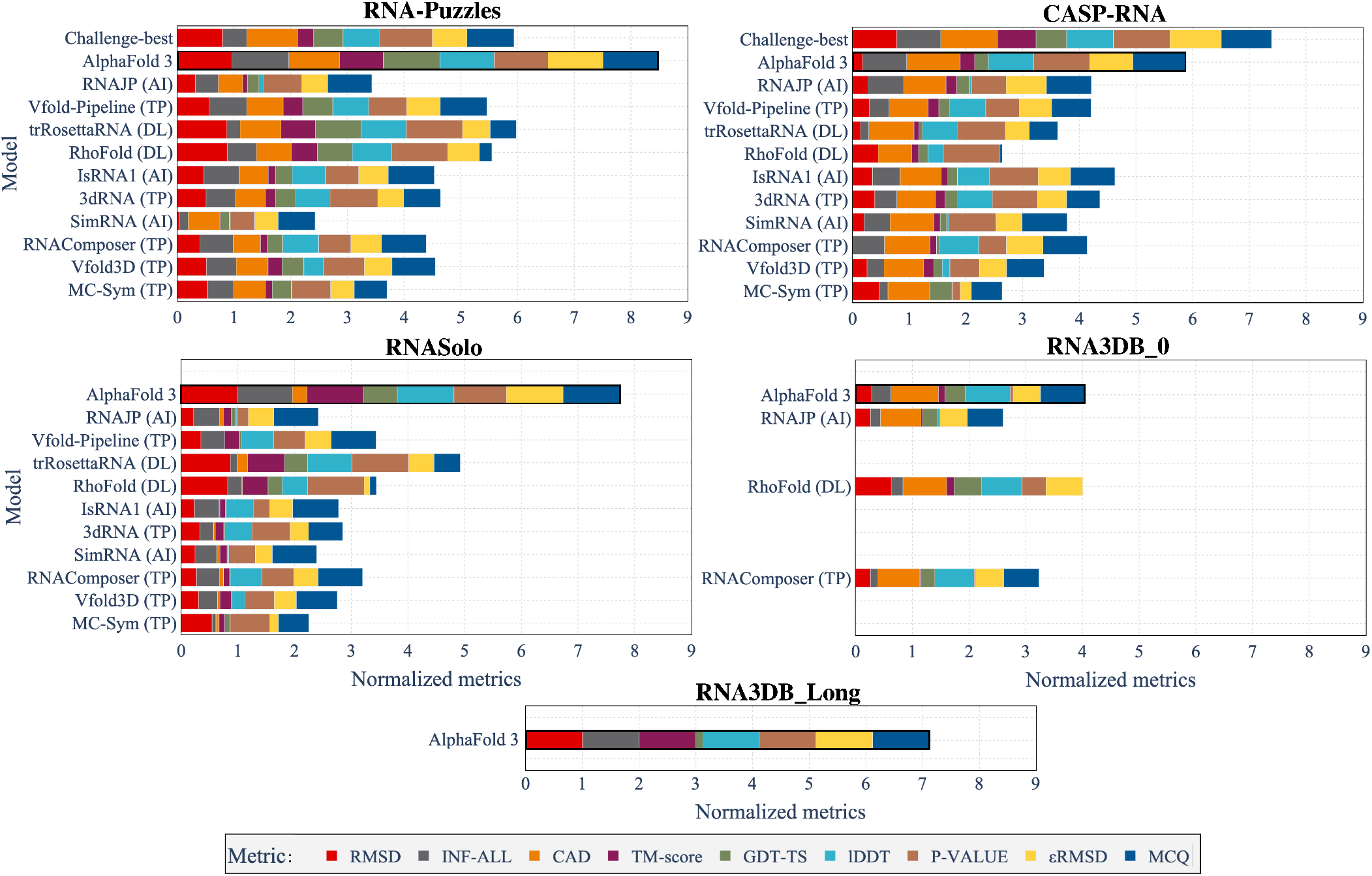
Cumulative normalised metrics for each of the benchmarked methods (2, 4, 6, 12, 16–18, 25, 28, 32, 58) for our five test sets. Each metric is normalised by the maximum value over the five test sets, and the decreased metrics are inversed to have better values close to 1. *Challenge-best* means the best solutions from the RNA-Puzzles and CASP-RNA competitions. Types of methods are also mentioned with the names: DL for deep learning, TP for template-based and AI for *ab initio*. Methods are sorted by release time (except for *challenge-best*).

The best models from the CASP-RNA competitions, which are human-guided, outperform AlphaFold 3 for every metric for the CASP-RNA dataset. On the other hand, AlphaFold 3 shows a cumulative sum of metrics greater than the other methods for the other test sets. For RNA-Puzzles, the challenge-best solutions are from older solutions, with less advanced architectures compared to the more recent CASP-RNA solutions. For the RNA3DB_0 dataset, AlphaFold 3 performances are almost similar to RhoFold, which has a higher RMSD but a lower MCQ. AlphaFold 3 always has a high MCQ value, indicating it returns structures which are more physically plausible than *ab initio* methods (that use physics properties in their predictions). Nonetheless, it does not always have the best RMSD (outperformed in CASP-RNA and RNA3DB_0), suggesting that AlphaFold 3 does not always have the best alignment (in terms of all atoms) compared to the reference structure.

To compare the global performances of each type of approach, we report in Figure 5 the averaged metrics over the different types of approaches depending on the sequence length. We grouped the results for structures with a sequence length window of 25 nt (each point represents the mean computed on the best results per approach with sequence length from this 25 nucleotide window). Results on the other metrics are shown in Figure S1 in the Supplementary file.

**Figure 5.**
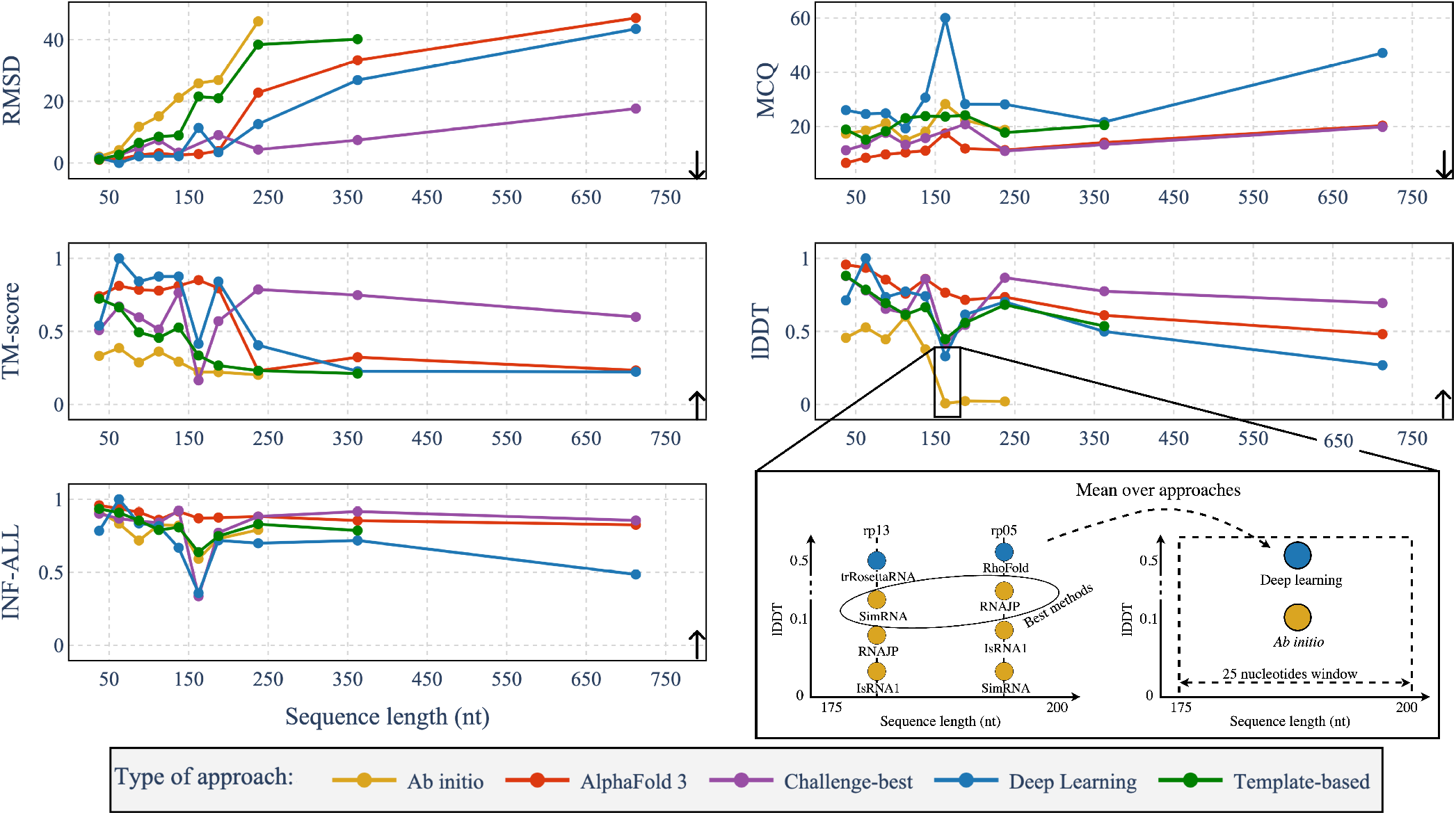
Averaged metrics depending on the sequence length for the different approaches (AlphaFold 3, *ab initio*, deep-learning, template-based and *challenge-best*). Each point represents the metric averaged over the best models of each approach for a window of 25 nt, from 25 to 750 nt. *Ab initio* methods group RNAJP (2), IsRNA1 (4) and SimRNA (6) while template-based methods group Vfold-Pipeline (12), 3dRNA (16), RNAComposer (17), Vfold3D (18) and MC-Sym (58). Deep learning methods group trRosettaRNA (28) and RhoFold (25). Metrics are computed for the RNA-Puzzles, CASP-RNA and RNASolo datasets. The *Challenge-best* corresponds to the best results from either RNA-Puzzles or CASP-RNA competitions but does not appear for the RNASolo dataset. The metrics are RMSD, MCQ (68), TM-score (62, 69), lDDT (66) and INF-ALL (37). RMSD and MCQ are descending (down arrow), i.e. the lower, the better, while TM-score, lDDT and INF-ALL are ascending (up arrow), i.e. the higher, the better.

All the benchmarked *ab initio* methods could not predict structures with sequences higher than 200 nt (or at least not with web servers). The best results from CASP-RNA and RNA-Puzzles challenges outperform AlphaFold 3 for every metric, except for structures with sequences between 150 and 250 nt. It seems to tend to increase with the sequence length for all kinds of approaches for the RMSD and MCQ and decrease for lDDT and TM-score. The results on INF-ALL show overall good results with values higher than 0.5 for almost every method. We also observe systematically higher values of MCQ for deep learning approaches compared to other methods. AlphaFold 3 seems to have predictions with better MCQ scores compared to other approaches, outperforming in terms of MCQ the challenge best solutions for structures with sequences below 200 nt.

These results show that AlphaFold 3 has competitive results without being the best model yet for all the challenges. It has behaviours different from existing deep learning approaches with much higher MCQ and, thus, torsional angles.

### The performance of AlphaFold 3 relative to sequence length

As seen previously, the prediction of RNA 3D structures usually becomes harder when the sequence length increases. Indeed, the *ab initio* methods fail to predict long interactions as the computation time highly increases with the sequence length. The template-based approaches are limited by the small number of long RNAs, as well as the deep learning methods, as shown in (30).

To observe more in detail the relation between sequence length and AlphaFold 3 performances, we report in Figure 6 the RMSD, MCQ, TM-score, lDDT and INF-ALL metrics depending on sequence length (for the five test sets). Links between the other metrics and the sequence length are available in Figure S2 of the Supplementary file.

**Figure 6.**
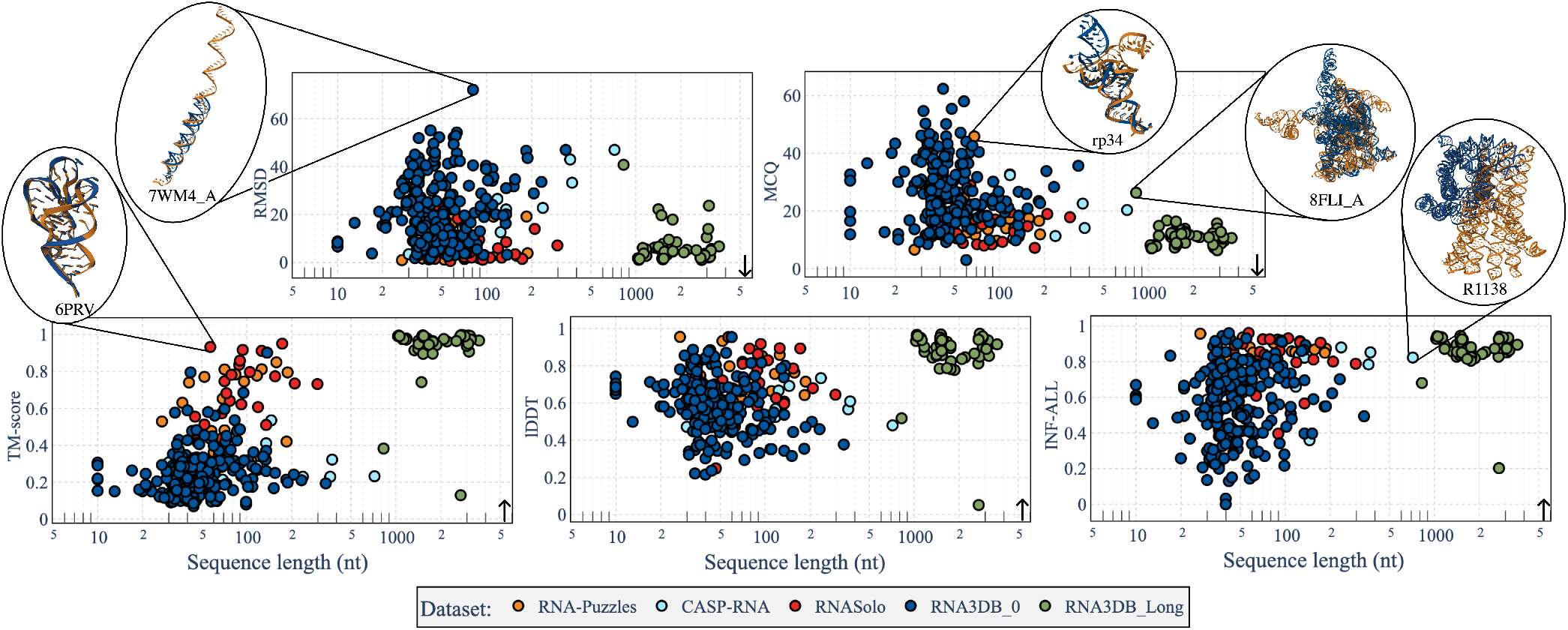
Dependence of metrics with the sequence length on the prediction of AlphaFold 3 (32) on the five test sets. For some of the predictions, we show the predicted structure (in blue) aligned with the native one (in orange) using US-Align (69). Metrics are RMSD, MCQ (68), TM-score (62, 69), lDDT (66) and INF-ALL (37). RMSD and MCQ are descending (down arrow), i.e. the lower, the better. TM-score, lDDT and INF-ALL are ascending (up arrow), i.e. the higher, the better.

Figure 6 indicates that, except for the RNA3DB_0 dataset, the RMSD increases for sequences between 0 and 1000 nt. For the RNA3DB_Long dataset with sequences higher than 1000 nt, the predictions have good results for every metric. We also observe a tendency of decrease for the lDDT, TM-score and INF-ALL (smaller decrease) when the structures have sequences higher than 100 nt (and below 1000 nt). For every metric, the predictions for the RNA3DB_0 dataset seem to have no clear dependence on the sequence length. For the other test sets with structures with sequences between 200 and 1000 nt, there is a common tendency of decrease in terms of performances for the AlphaFold 3 predictions.

### AlphaFold 3 results on long RNAs

Current methods for the prediction of RNA 3D structures are limited for long RNAs and hardly predict structures with sequences longer than 200 nt. AlphaFold 3 is, to the best of our knowledge, the only method that can predict long RNAs (with sequences higher than 1000 nt). Its predictions on RNA3DB_Long are of good performance, as shown in Figure 4. The only metrics where the results are not good are the GDT-TS and the CAD-score, which might be due to an error in computation.

The good results for long RNAs can be explained by the types of structures used in RNA3DB_Long. Indeed, all of the structures (except for one) are ribosomal RNAs, and thus have a lot of redundancy. This might be reflected in the PDB, which has been memorised by AlphaFold 3 during its training. As AlphaFold 3 uses the MSA as inputs, it could find similarities with trained structures and thus return excellent predictions if the families are well known. Most of the long RNAs in the PDB share common structures in the ribosomal family. Therefore, these results show a good generalisation of already-seen families from AlphaFold 3.

We report the two worst predictions of AlphaFold 3 on the RNA3DB_Long dataset in Figure 7. The two worst predictions for the other test sets are provided in Figure S3 of the Supplementary file. The RMSD for the two structures is relatively high (superior to 23Å). The second worst structure has a high TM-score (0.9), meaning that even for a long structure (3096 nt), the global alignment of atoms is well predicted. The INF-ALL is also high for these structures (higher than 0.68), meaning it returns a high proportion of key RNA interactions.

**Figure 7.**
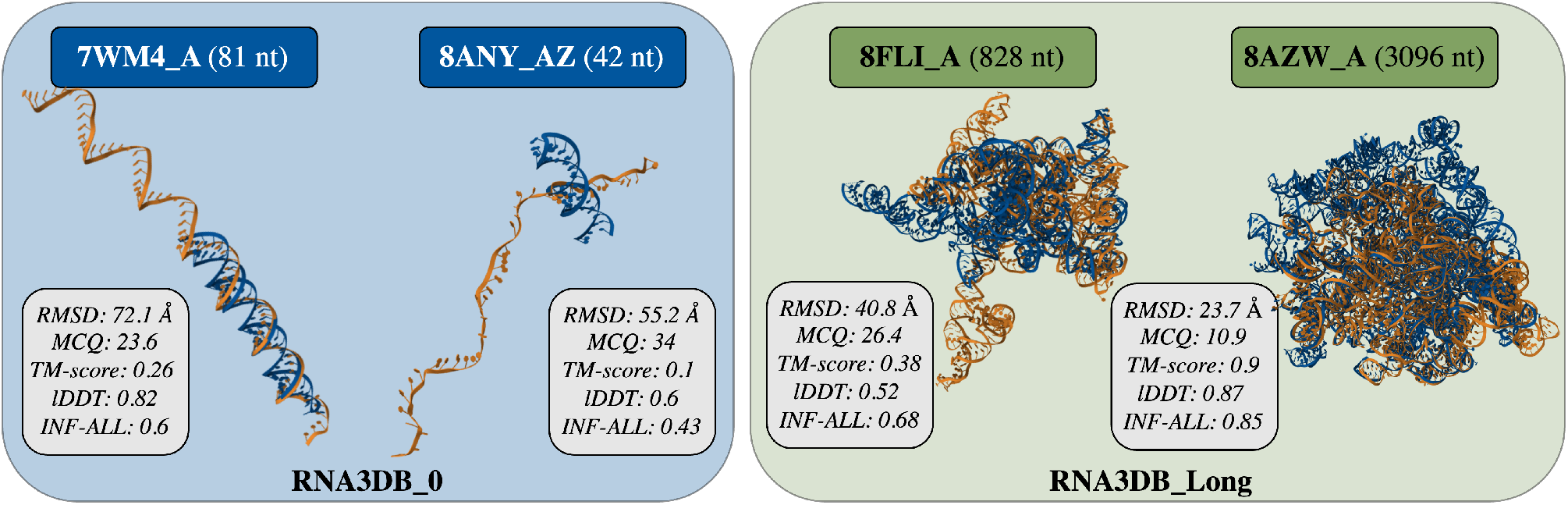
Worst two predicted structures from AlphaFold 3 (32) for RNA3DB_0 (left) and RNA3DB_Long (right) datasets. The RMSD, MCQ (68), TM-score (62, 69), lDDT (66) and INF-ALL(37) are provided for each structure. The predictions from AlphaFold 3 (in blue) are aligned with the native ones (in orange) using US-align (69).

### AlphaFold 3 results on orphan structures

The RNA3DB_0 dataset is mainly composed of structures without any hit in the Rfam family, which constitutes or- phan structures. The results of AlphaFold 3 for this dataset, as presented in Figure 4 and Figure 6, show overall lower performances compared to the other datasets. AlphaFold 3 achieves similar results compared to RhoFold for this dataset. These bad predictions for the RNA3DB_0 dataset could be explained by the non-common structures, as RNA3DB_0 has a lot of orphan structures. AlphaFold 3 uses multiple sequence alignment (MSA) as one of its inputs and thus is not restricted by structures from unknown families.

We detail the two worst predictions for RNA3DB_0 from AlphaFold 3 in Figure 7. We observe bad results in terms of metrics (high RMSD and MCQ values and low TM-score and INF-ALL) for the two structures. These structures also have a small number of nt (81 and 42), meaning that AlphaFold 3 might not fail because of the long-range interactions. Instead, these structures do not have a known family that could help AlphaFold 3 generalize well. It shows the limitation of AlphaFold 3 on orphan structures, where the generalisation of a large-scale model is hardly applicable even with small structures.

### AlphaFold 3 results on key RNA interactions

To evaluate the ability of AlphaFold 3 to predict non-canonical interactions, we depict the scatter plots between non-Watson-Crick INF (INF-NWC) and Watson-Crick INF (INF-WC) in Figure 8. The size of the points is proportional to the RMSD of structures and, thus, to their global atoms alignment. We observe a tendency to have a low RMSD (small points) whenever the INF-WC and INF-NWC are high. There are also many structures with an INF-NWC of 0, suggesting that AlphaFold 3 does not predict any of the non-Watson-Crick interactions (mostly for the RNA3DB dataset). Examples of successful and missing non-Watson-Crick interactions are shown in the Figure. For the results on stacking interactions, there are predictions where AlphaFold 3 does not predict the Watson-Crick interactions well but still predicts the stacking ones. It can be explained by good skeleton predictions while lacking the base conformations that produce the WC interactions. Secondly, there is an increased tendency between the INF-STACK and INF-WC: when AlphaFold 3 predicts the WC interactions well, it also tends to estimate the stacking well. Indeed, the stacking interactions can be more easily inferred whenever the WC interactions are respected.

**Figure 8.**
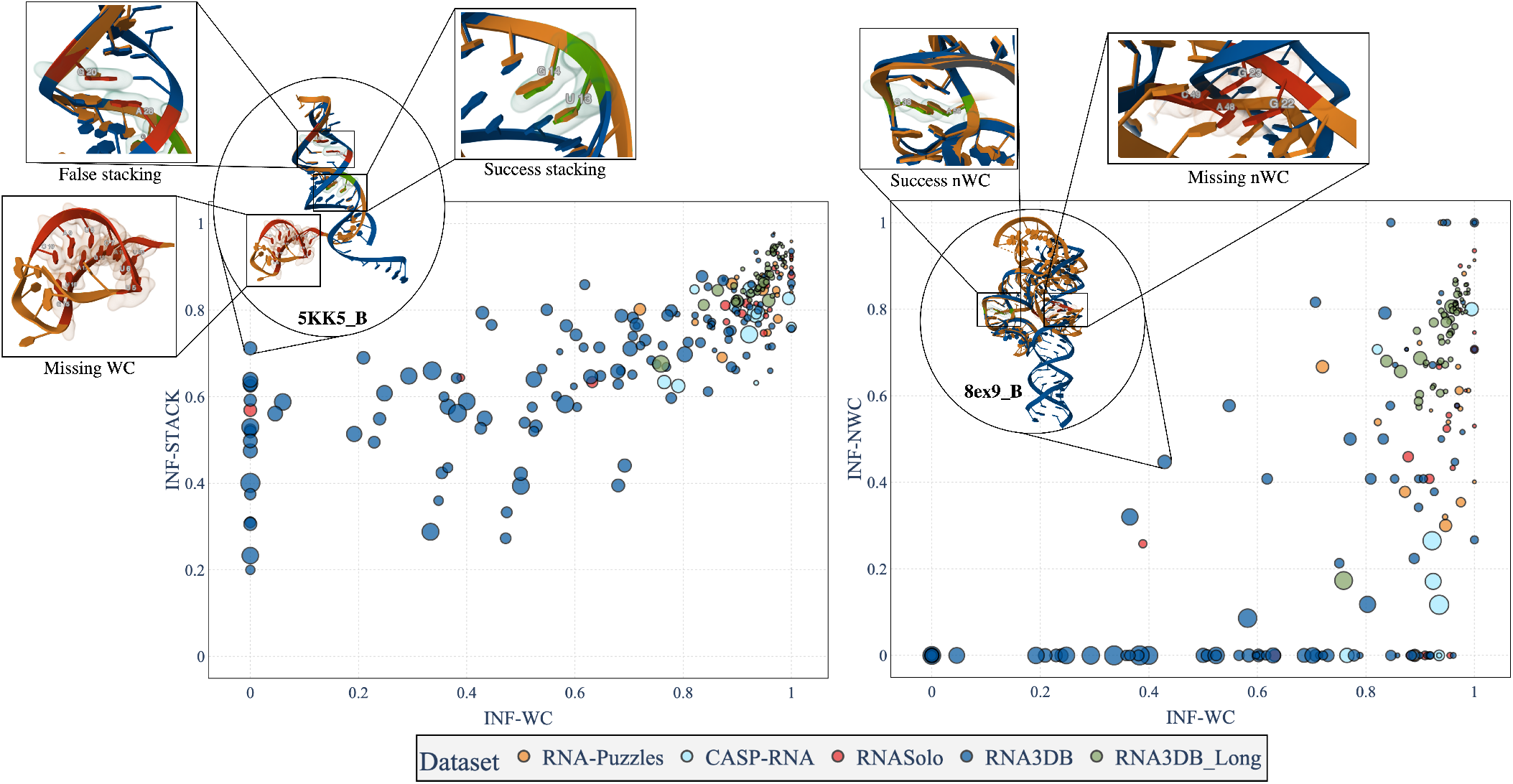
Link between INF Watson-Crick (WC) and non-Watson-Crick (nWC) and stacking (STACK) interactions for the predictions of AlphaFold 3 for our five test sets. The size of each point is proportional to the RMSD: the lower, the better. Only structures with at least one non-Watson-Crick interaction are shown in the figures. An INF (37) value of 1 means accurate reproduction of key RNA interactions, while a value near 0 means the structure does not reproduce the interactions. Left: INF non-Watson-Crick (INF-nWC) depending on INF Watson-Crick (INF-WC) interactions. Right: INF stacking (INF-STACK) depending on INF Watson-Crick (INF-WC) interactions.

To compare the key RNA interactions predicted from AlphaFold 3 with existing solutions, we present in Figure 9 the mean INF metrics (INF-WC, INF-NWC and INF-STACK) over RNA-Puzzles, CASP-RNA and RNASolo for the ten benchmarked models. Details for each dataset are provided in Table S1 of the Supplementary file. We show the results only on these datasets as we had complete predictions for each model only for these three datasets. AlphaFold 3 has better values for each INF metric compared to the other methods. The second best method to reproduce RNA key interactions is RNAComposer. While having good overall results in terms of cumulative metrics, trRosettaRNA shows bad results in terms of key RNA interactions. Even if AlphaFold 3 outperforms other solutions for all the INF metrics, the results for nWC interactions remain low (below 0.5), meaning there is still progress to reproduce RNA-specific interactions well.

**Figure 9.**
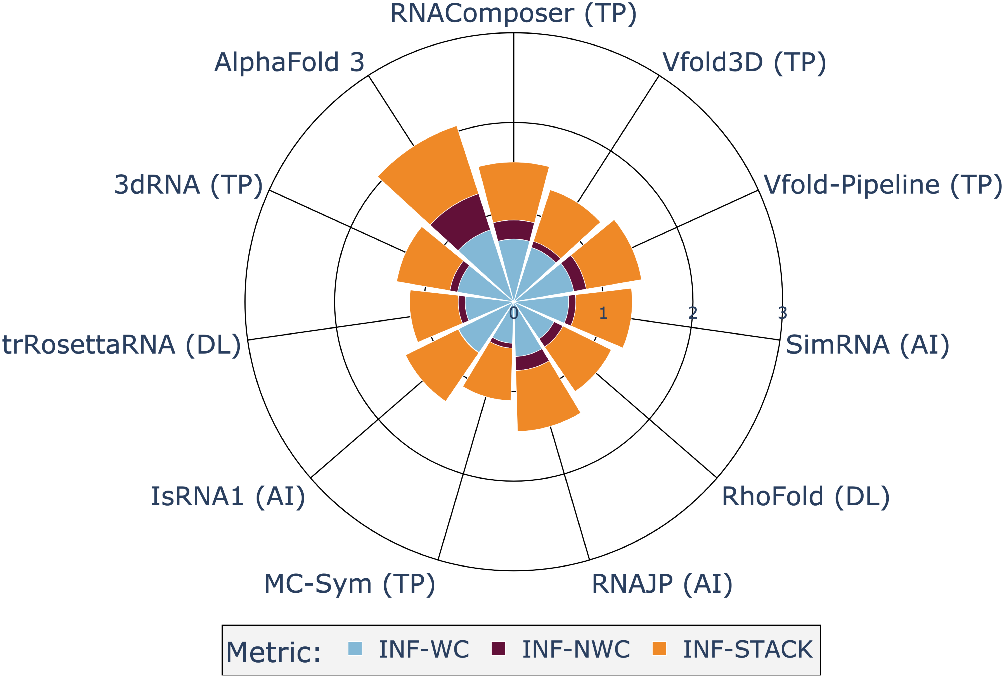
INF metrics for the different benchmarked models averaged over three test sets: RNA-Puzzles, CASP-RNA and RNASolo. INF metrics consider Watson-Crick (INF-WC), non-Watson-Crick (INF-NWC), and stacking (INF-STACK) interactions.

### Computation time

AlphaFold 3 is a deep learning method that has a complex architecture. Compared to existing *ab initio* methods, deep learning methods tend to be faster for inference. We report the computation time for a small RNA molecule (27 nt) and a long RNA (434 nt) for RNAComposer (17), RhoFold (25), trRosettaRNA (28), RNAJP (2) and AlphaFold 3 (32) in Table 1. We report the computation time of the fastest methods, while the time for the rest of the methods is available in our previous work (30). As we could only run RNAJP locally and each web server has different configurations, there is a bias in the comparison. RNAComposer, RhoFold and trRosettaRNA all predict small RNA (in less than a minute) very quickly, while RNAJP takes two hours (with default parameters). For a structure with a longer sequence, this is RNAComposer that has the fastest computation time (around three minutes). The *ab initio* method, RNAJP, takes 15 hours. AlphaFold 3 returns prediction in around five minutes, which shows fast inference. For RNA of very long sequences (around 3000 nt), AlphaFold 3 take multiple hours to predict (and sometimes returns errors and needs to be run multiple times to get results).

**Table 1.** Computation time for a sequence of 27 nt (PDB ID: 6Y0Y) and 434 nt (PDB ID: 7XD6). Computation time is computed using web servers except for RNAJP. Methods are sorted by release time. The types of approaches are either template-based (TP), *ab initio* (AI) or deep learning (DL).

| Model | Approach | Time<br>(min)<br>(27nt) | Time<br>(min)<br>(434nt) |
| --- | --- | --- | --- |
| RNAComposer (17) | TP | 1 | 3 |
| RhoFold (25) | DL | 1 | 10 |
| trRosettaRNA (28) | DL | 1 | 600 |
| RNAJP* (2) | AI | 120 | 900 |
| AlphaFold 3 (32) | DL | 2 | 5 |

\* RNAJP computation time is computed locally with a simulation time set to 50 × 10<sup>6</sup> steps on an NVIDIA P1000.

## Discussion

AlphaFold 3 is a deep learning method that has widened its scope to predict RNA structures (as well as other molecules) compared to its previous approach. Through our benchmark, we showed that AlphaFold 3 is a competitive method that outperforms most of the existing solutions. It yields better results for the CASP-RNA challenge and RNASolo, but remains outperformed by the best solutions from CASP-RNA challenge.

AlphaFold 3 has achieved good generalisation properties for the ribosomal structures (RNA3DB_Long dataset). This shows bias from existing data for RNA: most of the long RNA available in the PDB is ribosomal-related RNA.

AlphaFold 3 returns results with an overall good reproduction of RNA key interactions compared to existing solutions. It is also the best method to reproduce RNA torsional angles (best results in terms of MCQ), which was lacking in the existing deep learning methods (30).

There remain limitations that need to be addressed regarding the RNA folding problem. AlphaFold 3 does not reproduce all the non-Watson-Crick interactions, which is essential for the stability of RNA 3D structures. Furthermore, AlphaFold 3 fail to predict structures from orphan families (RNA3DB_0 dataset). These structures are hard to predict as there is no hint in the available data, which shows the limitation of generalisation to all the existing RNAs. AlphaFold 3, while reducing the impact of MSA on its architecture, still uses it, restricting its scope for RNA (as there are still unknown families). The computation time for the inference is very fast but remains limited by its usage in web servers (and the lack of available code).

## Conclusion

AlphaFold 2 had a huge success in the prediction of protein folding and has changed the field by the quality of its predictions. The new release of AlphaFold, named AlphaFold 3, has extended the model to predict all molecules from the PDB, like ions, ligands, DNA or RNA.

Through an extensive benchmark on five different test sets, we have evaluated the quality of predictions of AlphaFold 3 for RNA molecules. We have also compared the results with ten existing methods, which are easily reproducible as their predictions are available using web servers.

Our results show that AlphaFold 3 is of competitive quality, as it outperforms most of the existing solutions. It returns more physically plausible structures than *ab initio* methods. It outclasses existing deep-learning approaches for every dataset while better reproducing key RNA interactions and torsional angles. It also returns predictions very quickly compared to *ab initio* or current template-based approaches (but does not exceed RNAComposer (17) for inference time). For ribosomal long RNAs, AlphaFold 3 returns highly accurate predictions. It could be explained by its capability to generate structures from known families, which have been seen in its training data. As there is not a lot of data available, it is difficult to find complex structures without any homologous to evaluate performances fairly.

Nonetheless, AlphaFold 3 has not yet reached RNAs with the same success as proteins. Its new architecture allows the prediction of wide molecules but remains limited and hardly predicts non-Watson-Crick interactions. It does not generalize well on orphan structures, which are not related to any RNA-known families.

The prediction of atom coordinates instead of base frames, as done in AlphaFold 2, allows the extension of predictions for a wide range of molecules but prevents the generalisation of RNA-specific interactions. The lack of data is also a limitation that prevents the robustness of deep learning methods in general, and so is AlphaFold 3.

## ACKNOWLEDGEMENTS

This work is supported in part by UDOPIA-ANR-20-THIA-0013, Labex DigiCosme (project ANR11LABEX0045DIGICOSME), performed using HPC resources from GENCI/IDRIS (grant AD011014250), and operated by ANR as part of the program “Investissement d’Avenir” Idex ParisSaclay (ANR11IDEX000302).

