## Supplementary file for "Has AlphaFold 3 reached its success for RNAs?"

### Has AlphaFold 3 reached its success for RNAs? Supplementary file

July 19, 2024

#### Results

| | CASP-RNA | | | RNA-Puzzles | | | RNASolo | | |
|---|---|---|---|---|---|---|---|---|---|
| | WC | NWC | STACK | WC | NWC | STACK | WC | NWC | STACK |
| Challenge-best | **0.81** | **0.38** | 0.73 | 0.60 | 0.22 | 0.64 | - | - | - |
| AlphaFold 3 | **0.81** | 0.21 | **0.74** | **0.89** | **0.55** | **0.83** | **0.83** | **0.51** | **0.82** |
| RNAJP | 0.69 | 0.12 | 0.72 | 0.55 | 0.15 | 0.65 | 0.59 | 0.19 | 0.67 |
| Vfold-Pipeline | 0.63 | 0.00 | 0.56 | 0.78 | 0.29 | 0.70 | 0.62 | 0.14 | 0.62 |
| trRosettaRNA | 0.50 | 0.03 | 0.54 | 0.60 | 0.13 | 0.57 | 0.53 | 0.07 | 0.52 |
| RhoFold | 0.31 | 0.03 | 0.54 | 0.70 | 0.20 | 0.66 | 0.50 | 0.10 | 0.60 |
| IsRNA1 | 0.65 | 0.00 | 0.65 | 0.80 | 0.00 | 0.68 | 0.61 | 0.01 | 0.64 |
| 3dRNA | 0.61 | 0.02 | 0.61 | 0.73 | 0.17 | 0.64 | 0.57 | 0.07 | 0.56 |
| SimRNA | 0.59 | 0.10 | 0.65 | 0.68 | 0.06 | 0.59 | 0.57 | 0.08 | 0.65 |
| RNAComposer | 0.73 | 0.10 | 0.65 | 0.75 | 0.33 | 0.67 | 0.62 | 0.21 | 0.62 |
| Vfold3D | 0.57 | 0.02 | 0.58 | 0.73 | 0.13 | 0.65 | 0.61 | 0.06 | 0.59 |
| MC-Sym | 0.45 | 0.00 | 0.58 | 0.68 | 0.04 | 0.64 | 0.26 | 0.12 | 0.54 |

Table 1: INF [12] metrics (INF-WC, INF-NWC and INF-STACK) for each method for three datasets: CASP-RNA, RNA-Puzzles and RNASolo. The *Challenge-best* corresponds to the best results from the either RNA-Puzzles or CASP-RNA competitions. Methods are AlphaFold 3 [16], RNAJP [1], VFold-Pipeline [4], trRosettaRNA [9], RhoFold [10], IsRNA1 [2], 3dRNA [5], SimRNA [3], RNAComposer [6], Vfold3D [7] and MC-Sym [8]. Methods are sorted by release time (except for *challenge-best.* )

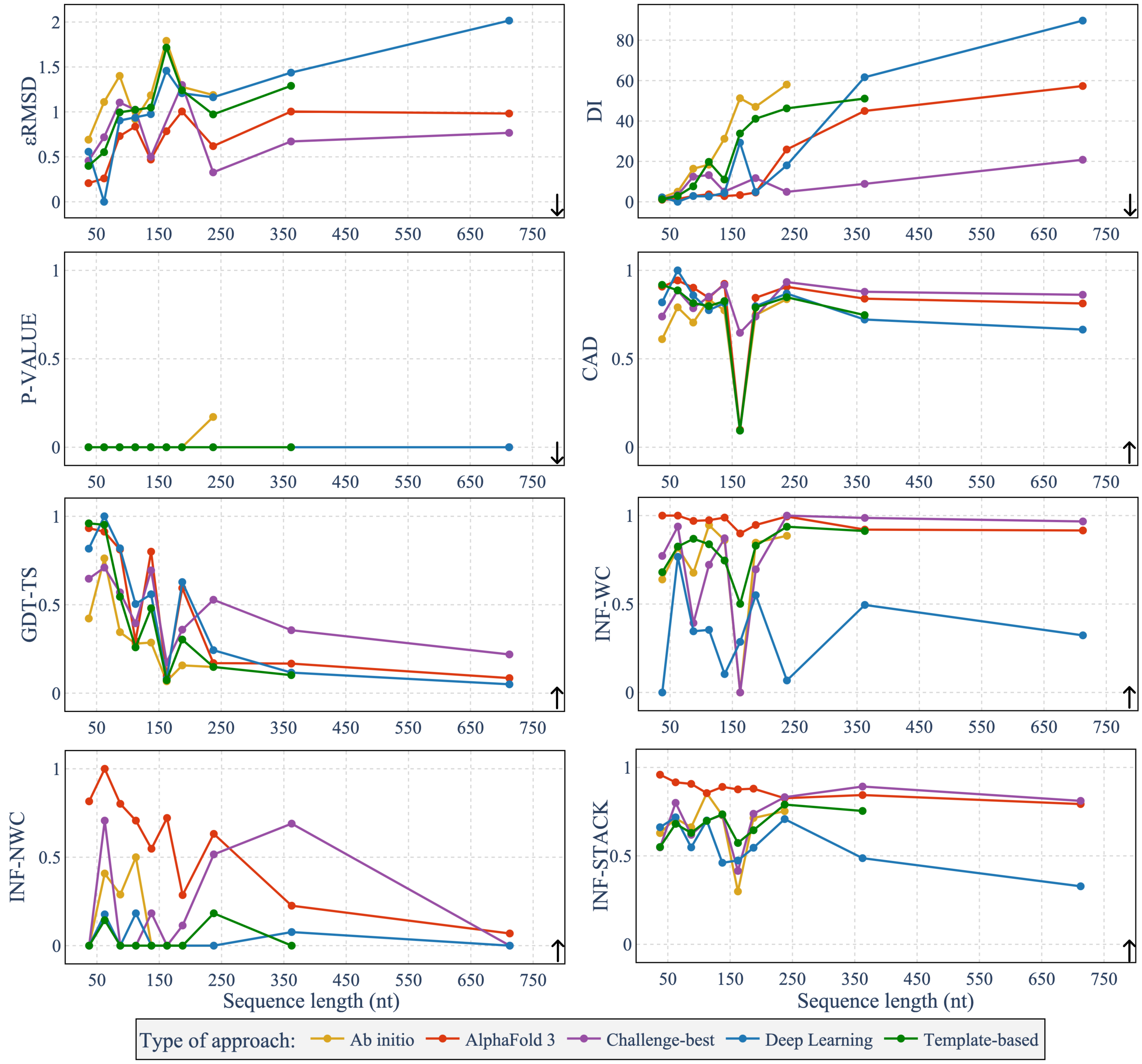


Figure S1: Averaged metrics depending on the sequence length for the different approaches (AlphaFold 3, *ab initio*, deep-learning, template-based and *challenge-best*). Each point represents the metric averaged over the best models of each approach for a window of 25 nucleotides, from 25 to 750 nucleotides. *Ab initio* methods group RNAJP [1], IsRNA1 [2] and SimRNA [3] while template-based methods group Vfold-Pipeline [4], 3dRNA [5], RNAComposer [6], Vfold3D [7] and MC-Sym [8]. Deep learning methods group trRosettaRNA [9] and RhoFold [10]. Metrics are computed for the RNA-Puzzles, CASP-RNA and RNASolo datasets. The *Challenge-best* corresponds to the best results from either RNA-Puzzles or CASP-RNA competitions but does not appear for the RNASolo dataset. The metrics are $\epsilon$RMSD [11], DI [12], P-VALUE [13], CAD-score [14], GDT-TS [15] and INF metrics [12] (INF-WC, INF-NWC and INF-STACK). $\epsilon$RMSD, DI and P-VALUE are descending (down arrow), i.e. the lower, the better, while CAD-score, GDT-TS, INF-WC, INF-NWC and INF-STACK are ascending (up arrow), i.e. the higher, the better.

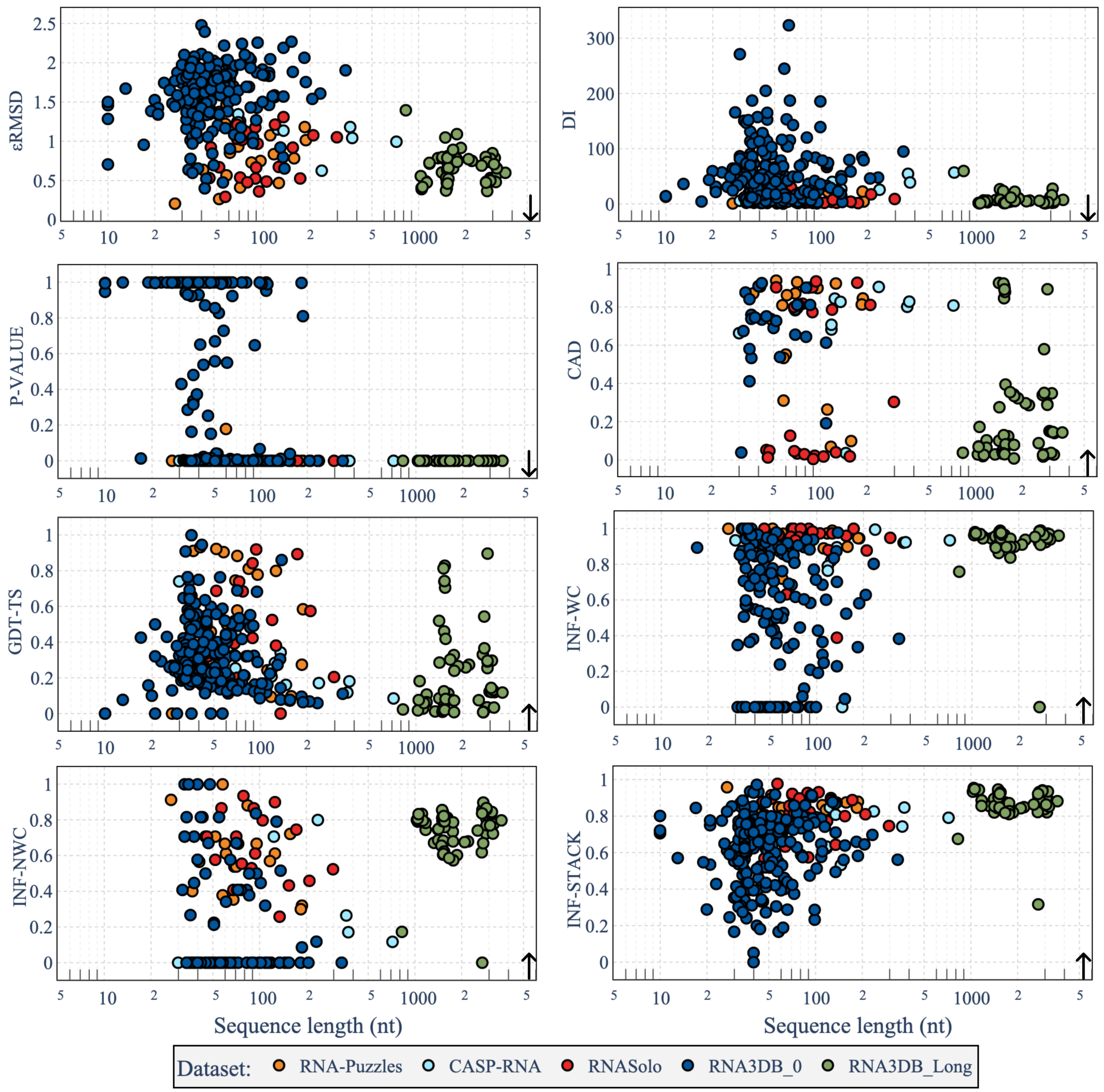


Figure S2: Dependence of metrics with the sequence length on the prediction of AlphaFold 3 [16] on the five test sets. Metrics are $\epsilon$RMSD [11], DI [12], P-VALUE [13], CAD-score [14], GDT-TS [15] and the INF [12] metrics (INF-WC, INF-NWC and INF-STACK). $\epsilon$RMSD, DI and P-VALUE are descending (down arrow), i.e. the lower, the better. CAD-score, GDT-TS, INF-WC, INF-NWC and INF-STACK are ascending (up arrow), i.e. the higher, the better.

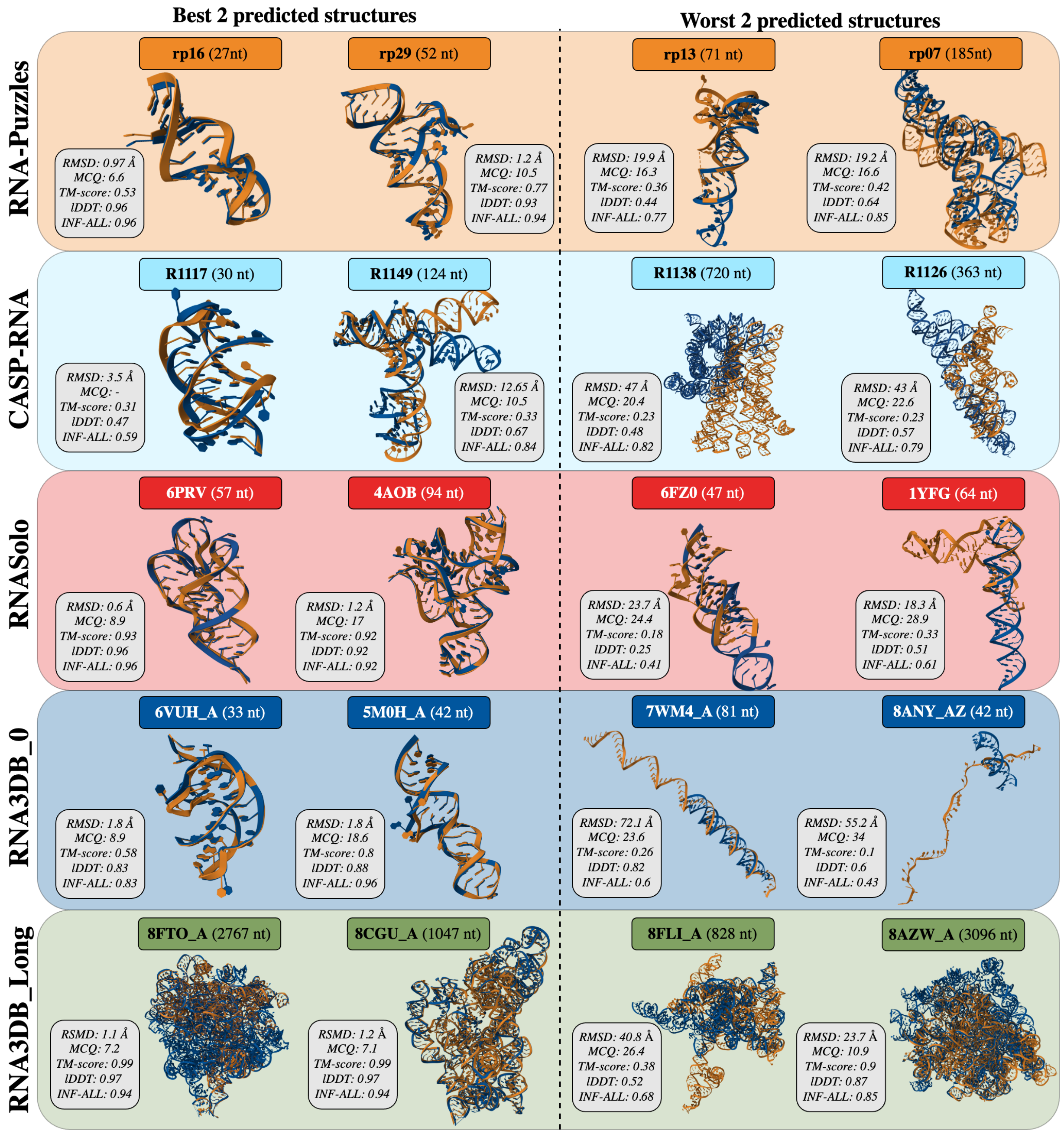


Figure S3: The two best and two worst predictions from AlphaFold 3 [16] for RNA-Puzzles, CASP-RNA RNASolo, RNA3DB_0 and RNA3DB_Long. The RMSD, MCQ [17], TM-score [18], lDDT [19] and INF-ALL [12] are included for each structure. The predictions from AlphaFold 3 (in blue) are aligned with the native ones (in orange) using US-align [20].
